# Structure and consistency of self-reported social contact networks in British secondary schools

**DOI:** 10.1101/322271

**Authors:** Adam J. Kucharski, Clare Wenham, Polly Brownlee, Lucie Racon, Natasha Widmer, Ken T. D. Eames, Andrew J. K. Conlan

## Abstract

Self-reported social mixing patterns are commonly used in mathematical models of infectious diseases. It is particularly important to quantify patterns for school-age children given their disproportionate role in transmission, but it remains unclear how the structure of such social interactions changes over time. By integrating data collection into a public engagement programme, we examined self-reported contact networks in year 7 groups in four UK secondary schools. We collected data from 460 unique participants across four rounds of data collection conducted between January and June 2015, with 7,315 identifiable contacts reported in total. Although individual-level contacts varied over the study period, we were able to obtain out-of-sample accuracies of more than 90% and F-scores of 0.49-0.84 when predicting the presence or absence of social contacts between specific individuals across rounds of data collection. Network properties such as clustering and number of communities were broadly consistent within schools between survey rounds, but varied significantly between schools. Networks were assortative according to gender, and to a lesser extent school class, with the estimated clustering coefficient larger among males in all surveyed co-educational schools. Our results demonstrate that it is feasible to collect longitudinal self-reported social contact data from school children and that key properties of these data are consistent between rounds of data collection.

## Introduction

Age-specific social mixing patterns are important in shaping the spread of infectious disease, from pandemic influenza [1, 2] to varicella and parvovirus [3]. As well as measuring contacts using proximity sensors such as radio-frequency identification (RFID) tags [4, 5, 6], self reported social contacts can be collected via routine questionnaires in different settings [7, 8]. Such data is commonly used to parameterise mathematical models of infectious diseases [9].

Mixing patterns of children are recognized as particularly important for understanding how disease spreads, with children often representing a key risk group [10, 11, 12]. They have limited acquired immunity to many pathogens, making them susceptible to infection [13, 3]. They also tend to make more social contacts than adults, and the majority of these social contacts are with other immunologically naive children [8], increasing the potential for transmission. As a primary location for children’s interactions, schools can therefore play an important role in the spread of infectious disease [1, 14, 15, 16, 17].

Surveys of social contacts depend on the subjective judgement of the participants and thus may be subject to recall and subconscious biases related to how the question is framed [18,19,20,21]. Although one study repeatedly measured contact networks among 49 university staff and students over 14 non consecutive days, [22], most previous studies of self-reported contact networks have typically surveyed individuals on a particular day and have not quantified how individuals’ reported contacts change over time [23, 24]. In particular, it has been difficult to establish how robust network structures are over time and how accurately these network structures represent regular interactions between children [25, 26]. A key concept in social network analysis is that social networks are dynamic and peer groups will change over time [27], yet this dynamic nature is rarely taken into consideration when considering how diseases spread between children.

To understand the shape and consistency of self-reported social interactions between children over time, we analysed social contact patterns in four secondary schools over a period of five months. Building on a previous project, which collected similar social contact data at a single point in time [23], we integrated data collection into a public engagement programme designed to teaching students how to conduct academic research and learn about disease dynamics, to quantify how year 7 students (aged 11-12) interacted within their classes and year-groups, as well as outside school. Further, we explored the robustness of self-reported contact surveys by repeating the same questionnaire at several points within the school year to identify how social networks changed over time.

## Methods

### Study design

In collaboration with the Millennium Mathematics Project, an educational and outreach project based at University of Cambridge, we recruited four schools to participate in our public engagement and research project in spring 2014. From the applications received, 4 schools were selected in a purposive sample to include a range of geographical and socioeconomic settings (north/south, rural/urban, single-sex/co-educational, fee paying/non-fee paying). For context, 83% of the UK population live in urban areas [28], 4% of schools in England are single-sex [29] and around 7% of children in Britain attend fee paying schools [30].

In the four schools, we worked with year 10 (i.e. 14-15 years old) students on the study design and logistics. Year 7 students (aged 11-12) within each school were the target population for data collection. As with previous projects [23, 31], we used public engagement with schools to facilitate data collection as part of a wider programme of curriculum-relevant scientific activities [32]. We did this using a combination of two school visits per school and six joint video conferences. These video conferences were particularly useful in enabling interaction between the four school classes involved in the project. In particular, all four schools collaborated to design the final questionnaire, obtain consent from study participants, data collection, and analysis and discussion of findings. This provided students with first hand experience of working on a research project from hypothesis to conclusion.

The final survey aimed to ascertain which people year 7 students spent the most time with on a given day. We used a process of peer nomination as a method for data collection [33]: students were asked, via the research questionnaire, to list the six other students in year 7 at their school that they spend the most time with. The choice of six for the number of named contacts follows from a previous study [23], which selected six to balance the risk of right-censoring (i.e. if students had more than six contacts) with the potential for deliberate over-reporting of contacts if there was no upper limit (i.e. as a result of ‘competitive naming’ via peer pressure). Students were also asked to report their gender, school class, as well as non-year group contacts. The specific question asked was ‘how many people not in your year group (including family and teachers) did you have a conversation with yesterday?’ The respondent could select one of five possible categories: 0-5 people; 6-10; 11-15; 16-20; 21 or more.

Collecting data in schools can be challenging for external researchers; we tried to mitigate these logistical constrains by having the students in year 10 administer paper questionnaires to year 7 students at regular intervals throughout the second half of the academic year 2014-15 (Table 1). This allowed the questionnaires to be carried out at suitable times based on student researcher’s knowledge of the school, but also attempted to mitigate external intervention in participant decision making. We encouraged the school participants to carry out the surveys at similar time periods (i.e. at monthly intervals) but there was some variation in timing of survey between schools, based on student availability linked to term times, exam periods, and timetabled registration sessions (Table 1).

**Table 1:**
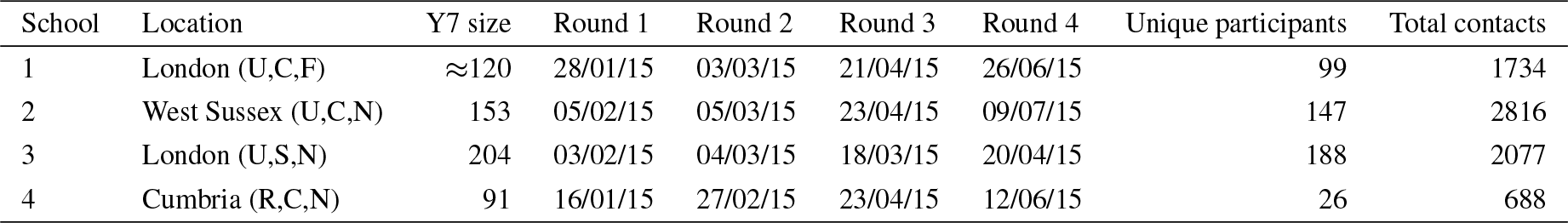
Summary of survey data from the four schools. Additional characteristics are given in parentheses: R=rural, U=urban; S=single-sex, C=co-educational; F=fee paying, N=non-fee paying. Unique participants denotes the number of uniquely identifiable students who completed the survey across all rounds. Total contacts gives the number of identifiable contacts reported across all participants.

**Table 2:** Variation in network statistics for each school. Clustering coefficients are shown for the full network, as well as for networks with males or females only. Community size estimates were obtained using edge betweenness (EB) and label propagation (LB). 10,000 bootstrap samples were used to obtain the estimates. Median values for each metric are shown, with 95% CIs in parentheses.

| Variable | School 1 | School 2 | School 3 | School 4 |
| --- | --- | --- | --- | --- |
| Mean participants | 77 | 121 | 92 | 24 |
| Diameter | 14 (10-21) | 11 (9-15) | 14 (11-20) | 7 (5-11) |
| Clustering coefficient | 0.4 (0.34-0.46) | 0.35 (0.31-0.39) | 0.18 (0.13-0.25) | 0.38 (0.24-0.51) |
| Clustering coefficient (male only) | 0.44 (0.33-0.54) | 0.43 (0.37-0.49) | 0.18 (0.13-0.25) | 0.6 (0-1) |
| Clustering coefficient (female only) | 0.41 (0.33-0.49) | 0.35 (0.3-0.4) | – | 0.37 (0.14-0.57) |
| Assortativity by gender | 0.92 (0.85-0.97) | 0.81 (0.74-0.87) | – | 0.48 (0.3-0.68) |
| Assortativity by class | 0.35 (0.27-0.44) | 0.12 (0.076-0.16) | 0.13 (0.071-0.2) | 0.11 (-0.033-0.24) |
| Communities (EB) | 11 (8-15) | 10 (7-13) | 11 (8-16) | 7 (5-9) |
| Communities (LP) | 14 (11-18) | 15 (12-18) | 14 (7-19) | 7 (5-10) |
| Participants per community (EB) | 7 (5.1-9.6) | 12 (9.3-17) | 8.4 (5.8-12) | 3.4 (2.7-4.8) |
| Participants per community (LP) | 5.5 (4.3-7) | 8.1 (6.7-10) | 6.6 (4.8-13) | 3.4 (2.4-4.8) |

Two types of informed consent were required for each of the year 7 participants completing the questionnaires. First, their parents or guardians were invited to sign a written consent form to allow their child to participate in the study. Second, the students themselves were asked to verbally consent to undertake the questionnaire each time. The project was approved by the LSHTM observational research ethics committee (ref 8769).

### Data cleaning and processing

Once the survey forms were completed, the data were digitised and cleaned (Supporting Data S1). We compiled a clean list of unique participants and contacts by using the stringdist R package [34] to match each reported name in the survey database to names in a year group list. We matched names using a ranking based on the Jaro-Winkler distances between first name and surnames in the two lists. This measures the minimum number of single-character transpositions needed to convert one word into the other. We ranked potential matches based on the combined distance between first name (*d*_1_) and surname (*d*_2_), with first names given a larger weighting: 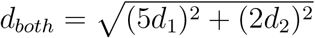. This criteria was obtained following several rounds of manual validation, with an additional round of validation performed after names were matched to identify erroneous matches. If *d*_*both*_ > 2, or the best match could not be validated manually, the participant or contact was not included in the analysis. Once the reported names had been cleaned and validated, all subsequent analysis was carried out on anonymised data, with each student name replaced by a unique code. Our analysis focused on two types of contacts. For each pair of participants in a specific round of data collection, a *single link* was defined if either one of the participants reported a contact between the pair (i.e. there was at least one unidirectional link, in either direction); a *mutual link* was defined if both participants reported contact with each other (i.e. a bi-directional link). Code and data required to reproduce all the analysis are available at: https://github.com/adamkucharski/schools-networks-15

### Network metrics

We used the igraph R package [35] to visualise the contact networks and examine five specific network metrics. These captured key aspects of network structure, as also examined in previous studies of contact patterns within schools [4, 23, 36]. The *global clustering coefficient*, which ranges from 0 to 1, is defined as the number of closed triplets in the network divided by number of connected trios of nodes, whether closed or not. Highly clustered networks have a coefficient value near one. The clustering coefficient for a specific round of data collection was calculated using all uni-directional links in the directed network generated from that round. The *diameter* of a network for a specific round was calculated from the shortest distance between the two nodes that are most distant, where the network included all uni-directional links. Given a categorical variable associated with each node, such as gender or school class, *nominal assortativity* measures how much nodes with the same categorical value are connected with one another [37]. Nominal assortativity, denoted *r*, is defined as:

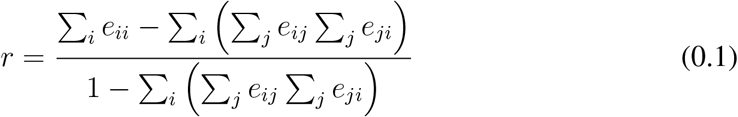

where *e*_*ij*_ is the proportion of edges that connect nodes of category *i* and *j*. Again, this was calculated using all uni-directional links in the network.

We also estimated how individuals were grouped within the network. We used two different community metrics. The first of these was the common benchmark method of *community detection based on edge betweenness*, which uses the Girvan-Newman algorithm [38]. This works by defining the ‘edge betweenness score’ for each edge in the network (i.e. the number of shortest paths between nodes in the network that pass through this edge), then the edge with the highest score is iteratively removed until a rooted tree remains, with each clade representing a community. As a sensitivity analysis, we also considered *community detection based on propagating labels* [39]. In this algorithm, a unique label is attributed to each node at the start, then each node iteratively adopts whichever label the majority of its neighbours have, until the labelling converges to identify communities.

### Bootstrap sampling

To assess the consistency of the network metrics for each school over time, we used bootstrap sampling [40]. In each iteration, we randomly sampled a subset *m* of participants from the full set of unique participants, where *m* is the mean number of participants across the four rounds (Table S1). For each sampled participant, we then randomly selected one of the rounds they participated in, and used this round as their reported contacts. This generated a bootstrap network from which we could calculate the relevant network statistics. The variation in these network statistics reflected the level of consistency in responses between rounds. If student participation and responses were identical across rounds, we would have obtained exactly the same network in each bootstrap sample. We performed 10,000 iterations of bootstrap sampling to estimate the median and 95% confidence interval for each network metric.

### Out-of-sample prediction

Collecting social contact data can be extremely time consuming. It would therefore be useful to establish whether multiple rounds of data collection generate a more consistent estimate of which individuals are linked. To examine how closely networks based on data collected in each round of surveys could predict the networks generated from data in subsequent rounds, we used an out-of-sample prediction approach. We focused on individuals who had participated in all four rounds of data collection. For each round, we calculated whether each possible pair of individuals were connected by a single link (i.e. at least one uni-directional link) in that round. We analysed single links rather than mutual links because we had already heavily thinned the networks by focusing only on participants who appeared in all rounds. We generated predictions using networks constructed from one or more rounds of training data; if a single link was present between two individuals in the training dataset, we predicted it would exist in later rounds. We then compared our predictions with test datasets, i.e. networks generated from later rounds of data collection. In the case where there was disagreement in the training datasets (e.g. a single link was present in two rounds of training data, but absent in the third), we defined the probability of this pair being connected by a single according to the proportion of training rounds that included a link. This make it possible to calculate expected numbers of true positives (TP) and true negatives (TN) and false positives (FP) and false negatives (FN). Specifically, these numbers were defined as:

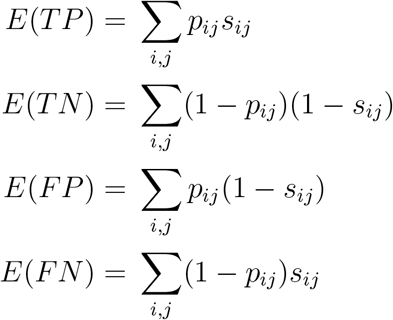

where *p*_*ij*_ is the proportion of training sets that have a single link between *i* and *j*, and *S*_*ij*_ = 1 if there is a single link between *i* and *j* in the test network. For example, in the situation where a single link is present in 2/3 of the training rounds, it would predict a single link in the test data with probability 2/3; if the single link had been present in all training rounds, it would have probability 1 of being predicted in the test data. Using the expected values of TP, TN, FP and FN, we calculated precision 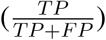, recall 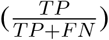, accuracy 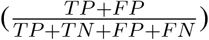 and F-score 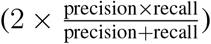.

As well as comparing network links, we measured the consistency of individual-level indegree across rounds. For each participant, we calculated their indegree in the training round, compared these predictions with the test data, then calculated the mean difference across all participants. If there were multiple training rounds, the mean was used. Suppose two rounds of training data were used and a participant had indegree 3 and indegree 4 in the training rounds and indegree 6 in the test round. In this case, the absolute difference was calculated to be |3.5 − 6| = 2.5). We used the same approach to calculate the mean difference in reported contacts outside the year group between training and test rounds.

## Results

In total, 460 unique year 7 participants completed 1,254 surveys over four rounds of data collection between January and June 2015, with 7,315 contacts reported in total (Table 1). In three schools, over 80% of the year group participated in at least one round of data collection; one school had considerably less participants in the dataset, the result of a low response rate in parental consent. Plotting all uni-directional links reported in each round of data collection, we found noticeable variation in contact links and active participants over time (Figure 1). In all rounds, however, we observed a clear gender divide in the year groups in the three co-educational schools (the other school was single-sex).

**Figure 1:**
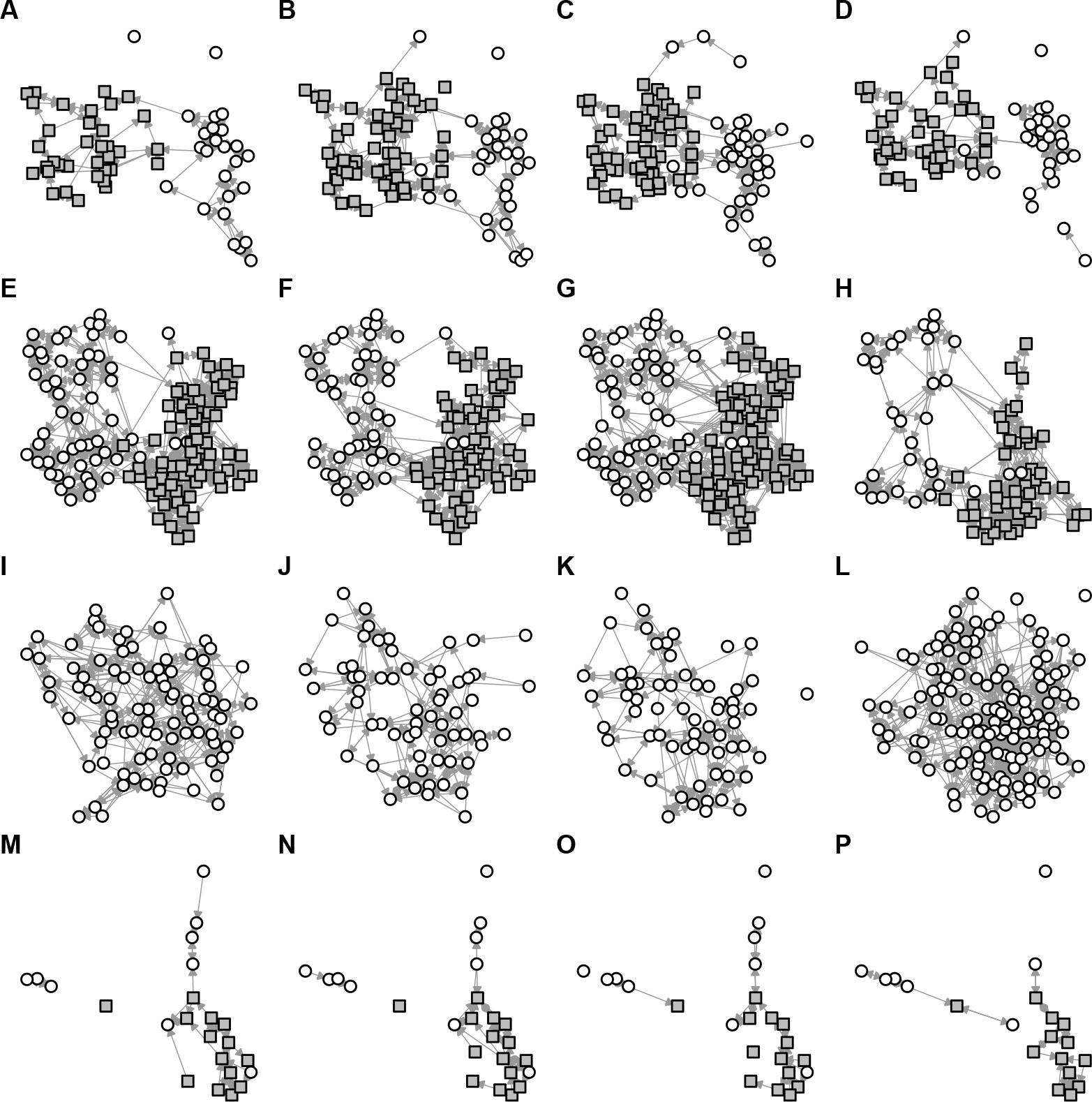
Reported social contacts in four schools over four months. (A-D) Coeducational school in London. All uni-directional links between study participants are shown, with nodes fixed in same position if participants were surveyed in multiple rounds and each arrow pointing toward the participant who was named. Grey squares show females, white circles show males. (E-H) Co-educational school in West Sussex. (I-L) Single-sex school in London. (M-P) Co-educational school in Cumbria.

We used several metrics to assess consistency in network structure over time. First we calculated the indegree distributions for each set of networks, and found that these followed a qualitatively similar pattern over time for each school, with considerable variation between schools (Figures 2A-D). Despite individual-level variation in contact reporting between surveys, network clustering within each school, as measured by the clustering coefficient, was relatively consistent over time, with estimates of 0.4 (95% bootstrap CI: 0. 34-0.46) for school 1, 0.35 (0.31-0.39) for school 2, and 0.18 (0.13-0.25) for school 3 (Table S1). School 3 had a significantly lower clustering coefficient than the other schools, suggesting less localised network structure. This reflects school timetabling: in this school, all students were set by ability in every subject, and set composition varied widely depending on the subject. Uncertainty was greater for school 4, which had low numbers of participants; the clustering coefficient was estimated to be 0.38 (0.24-0.51). When we calculated clustering coefficients for the other school networks with either males or females only included, we obtained larger clustering coefficients for males in all coeducational schools, with a statistically significant difference for school 2, which had the most participants (Table S1). To examine connectivity across networks consisting of all single links, we calculated the diameter of each network, which measures the shortest distance between the two nodes that are most distant. This varied across schools and across rounds (Table S1). In schools 1-3, which had most of the year group participating, median network diameter ranged between 11 and 14.

**Figure 2:**
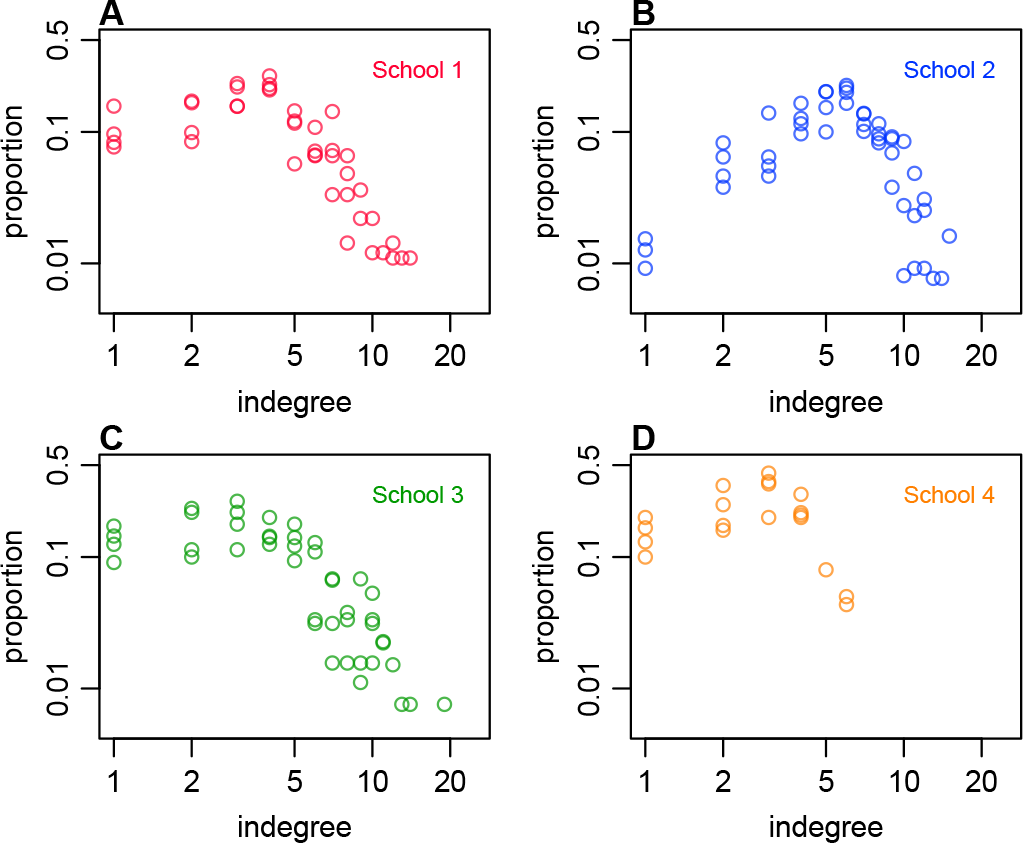
Distribution of contacts and clustering coefficient in each school, based on all single links in each of the four rounds of data collection. (A) Co-educational school in London. (B) Co-educational school in West Sussex. (C) Single-sex school in London. (D) Co-educational school in Cumbria.

Based on network assortativity, we found strong evidence of gender associated contact patterns in schools 1 and 2 (Table S1), as observed qualitatively in Figure 1. It was not possible to examine gender assortativity for school 3, which was single-sex. For school 1, there was also moderate assortativity based on the school class individuals were in, with positive but lower assortativity in schools 2 and 3. These associations are visible when single links from all four rounds of data collection are plotted together (Figure 3), and particularly prominent when only mutual links are plotted (Figure 3B). In addition, the largest components for schools 1 and 2 were dominated by females, whereas males tended to form smaller, less connected social groupings when mutual links were considered. Further, we used two community detection methods - edge betweenness and label propagation - to measure how many subgroups exist within the networks. The estimates based on the edge betweenness method varied between rounds suggested 7-12 communities within a year group for the three schools with high participation rates (Table S1); the label propagation method suggested the existence of 14-15 communities. Based on the numbers of participants in each school, this suggests each community contains 6-8 individuals.

**Figure 3:**
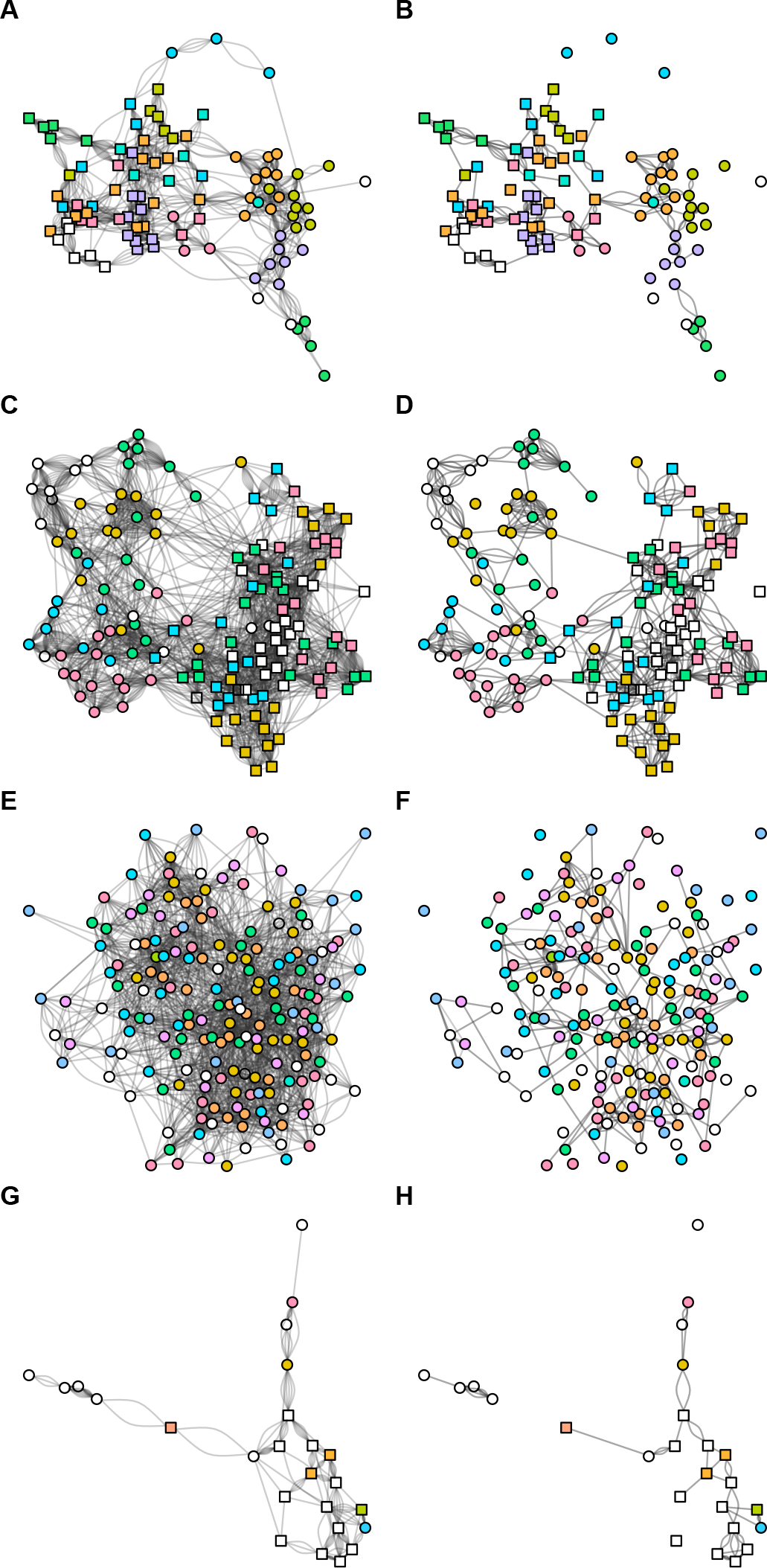
Structure of social contacts in the four schools across all sampling periods. (A-B) Co-educational school in London. Reported school classes are grouped by colour (white indicates no data); squares show females, circles show males. Left column shows all reported uni-direction links across all four rounds (i.e. maximum of eight possible links between each pair); right column shows only mutually reported (i.e. bi-directional) links across the found rounds (i.e. maximum of four possible links). (C-D) Co-educational school in West Sussex. (E-F) Single-sex school in London. (G-H) Co-educational school in Cumbria.

Because social contact surveys can require considerable effort to conduct, we also assessed the predictive power of single or multiple rounds of data collection. Using data only for participants that were present in all four rounds of data collection, we constructed networks of single links using either the first one, two or three consecutive rounds of data (as specified in Table 1), then compared these networks with the remaining test dataset(s). We assessed predictions about whether a single link existed between two specific participants by calculating the number of true and false positives and negatives (Table 3). We then calculated the precision, recall, F-score and accuracy of our out-of-sample predictions. We found that overall prediction accuracies ranged between 90-98%, but did not improve with the number of rounds of training data used (Figure 4A). Corresponding F-scores ranged between 0.49-0.84; again, these did not improve with the number of rounds of training data used (Figure 4B).

**Figure 4:**
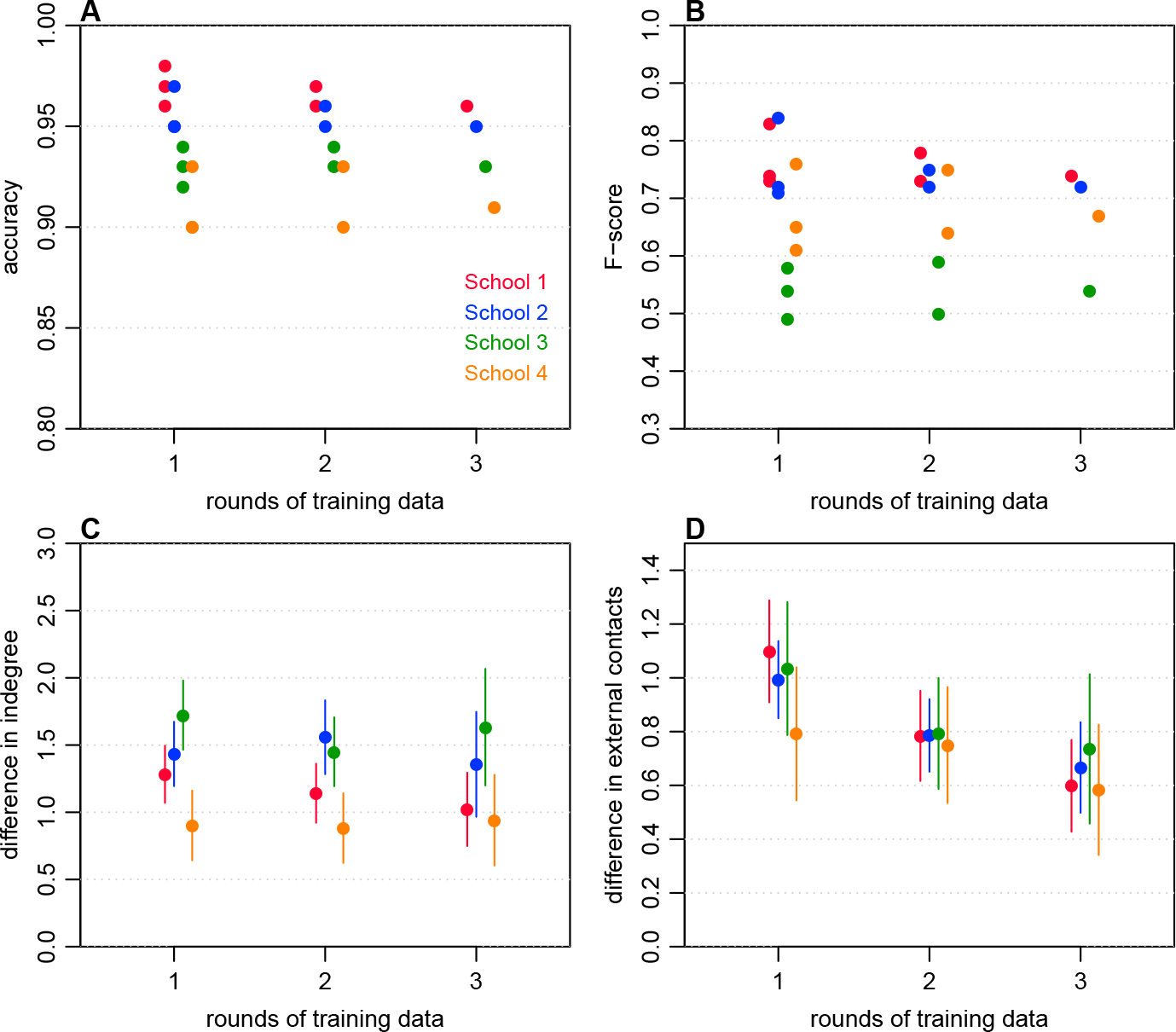
Predictive power of contact surveys. (A) Accuracy of predicting single links in the network when one, two or three rounds of data collection are used to make out-of-sample predictions, corresponding to values shown in Table 3. (B) F-scores when different numbers of rounds of data collection are used to make out-of-sample predictions (C) Mean absolute difference in degree across all participants. (D) Consistency of individual-level categorical responses (i.e. mean absolute difference between value of category reported in training rounds and reported in test round). Question was ‘How many people not in your year group (including family and teachers) did you have a conversation with yesterday?’ (1: 0-5, 2: 6-10, 3: 11-15, 4: 16-20, 5: 20+).

**Table 3:** Predictive power of surveys across rounds. Table shows which round(s) of data were used to train the network (i.e. identify which students are connected by a single link and not), and which round of data this network was tested against. ‘No link’ and ‘link’ indicates how many students are connected by a single link in the training data. When two training sets are used, students are sometimes linked in one and not linked in the other; these are classed as ‘ambiguous’. Potential links gives the total number of possible single links that could exist between nodes in that network. The ability of a training dataset to predict links in a test dataset is assessed using precision, recall, F-score and accuracy (see Methods).

| School | Train | Test | No link | Ambiguous | Link | Precision | Recall | F-score | Accuracy | Potential links |
| --- | --- | --- | --- | --- | --- | --- | --- | --- | --- | --- |
| 1 | 1 | 2 | 962 | 0 | 73 | 0.82 | 0.85 | 0.83 | 0.98 | 1035 |
| 1 | 1 | 3 | 962 | 0 | 73 | 0.71 | 0.78 | 0.74 | 0.97 | 1035 |
| 1 | 1 | 4 | 962 | 0 | 73 | 0.73 | 0.74 | 0.73 | 0.96 | 1035 |
| 1 | 2 | 3 | 951 | 24 | 60 | 0.75 | 0.81 | 0.78 | 0.97 | 1035 |
| 1 | 2 | 4 | 951 | 24 | 60 | 0.73 | 0.73 | 0.73 | 0.96 | 1035 |
| 1 | 3 | 4 | 943 | 0 | 49 | 0.74 | 0.73 | 0.74 | 0.96 | 1035 |
| 2 | 1 | 2 | 677 | 0 | 64 | 0.84 | 0.83 | 0.84 | 0.97 | 741 |
| 2 | 1 | 3 | 677 | 0 | 64 | 0.72 | 0.73 | 0.72 | 0.95 | 741 |
| 2 | 1 | 4 | 677 | 0 | 64 | 0.73 | 0.68 | 0.71 | 0.95 | 741 |
| 2 | 2 | 3 | 666 | 21 | 54 | 0.74 | 0.75 | 0.75 | 0.96 | 741 |
| 2 | 2 | 4 | 666 | 21 | 54 | 0.74 | 0.7 | 0.72 | 0.95 | 741 |
| 2 | 3 | 4 | 653 | 0 | 45 | 0.75 | 0.7 | 0.72 | 0.95 | 741 |
| 3 | 1 | 2 | 324 | 0 | 27 | 0.48 | 0.5 | 0.49 | 0.92 | 351 |
| 3 | 1 | 3 | 324 | 0 | 27 | 0.52 | 0.67 | 0.58 | 0.94 | 351 |
| 3 | 1 | 4 | 324 | 0 | 27 | 0.52 | 0.56 | 0.54 | 0.93 | 351 |
| 3 | 2 | 3 | 311 | 27 | 13 | 0.53 | 0.67 | 0.59 | 0.94 | 351 |
| 3 | 2 | 4 | 311 | 27 | 13 | 0.49 | 0.52 | 0.5 | 0.93 | 351 |
| 3 | 3 | 4 | 307 | 0 | 11 | 0.54 | 0.53 | 0.54 | 0.93 | 351 |
| 4 | 1 | 2 | 117 | 0 | 19 | 0.58 | 0.65 | 0.61 | 0.9 | 136 |
| 4 | 1 | 3 | 117 | 0 | 19 | 0.74 | 0.78 | 0.76 | 0.93 | 136 |
| 4 | 1 | 4 | 117 | 0 | 19 | 0.63 | 0.67 | 0.65 | 0.9 | 136 |
| 4 | 2 | 3 | 111 | 14 | 11 | 0.75 | 0.75 | 0.75 | 0.93 | 136 |
| 4 | 2 | 4 | 111 | 14 | 11 | 0.64 | 0.64 | 0.64 | 0.9 | 136 |
| 4 | 3 | 4 | 109 | 0 | 11 | 0.67 | 0.67 | 0.67 | 0.91 | 136 |

As well as considering pairwise single links, we also measured the mean individual indegree of each participant across the training rounds, and compared this with the indegree in a test data set. We used the same combinations of test and training data as for the network links (Table 3). Results were pooled based on the number of rounds included (i.e. 1, 2, or 3), and a mean and confidence interval estimated. Regardless of number of rounds used in the training data, the mean difference in indegree was in the range of 0.7-2 (Figure 4C). We also found no significant improvements in the predictions of mean indegree when more rounds were included in the training data. Finally, we examined reported contacts outside the school year-group, which were reported categorically in five bands (Figure 4D). Here, having multiple rounds of training data did significantly reduce prediction error, as measured by the mean absolute difference between the numbers of the categories reported, for two schools when two rounds of data collection were included rather than one. However, these differences were not significant when a Bonferroni correction was applied to account for multiple comparisons. Even with a single round of training data, the values were clustered near to 1, suggesting that participant responses were consistent enough to on average report contacts within two adjacent categories.

## Discussion

We assessed key features and structural characteristics of social networks in four UK secondary schools, and how these varied over time. As in previous studies [23, 31], the research was embedded within a public engagement project, with students and teachers in participating schools contributing to the study design and data collection. We found that although individual contacts within the year 7 group studied varied over the five month study period, out-of-sample prediction accuracy of contacts was over 90%. Further, many aspects of the overall network structure, such as extent of clustering and number of communities, remained relatively stable between rounds of data collection.

Children are key epidemiological group for many directly transmitted infectious diseases [10, 11, 12]. Our results show that it is feasible to collect longitudinal self-reported data from a group of year 7 children, and that data collected is reasonably consistent between rounds, which should increase confidence in cross-sectional studies examining self-reported contacts in this age group. We also observed considerable variation in network structure between the schools, indicating that while key network properties may be relatively robust within a school, there may be substantial differences depending on where a study is conducted. The small differences in individual-level indegree across rounds (Figure 4C) suggests that highly or weakly connected individuals remain so over time. However, we did not examine within or between classroom contact patterns in detail, given the low assortativity for classroom mixing within most schools. In future studies, it would be interesting to examine how the physical structure of buildings and timetabling influences social contacts [14], given that the school in our study that set students for all subjects had lower clustering than other schools.

There are some additional limitations to our study. To ensure the questionnaire was straightforward to complete, students were asked to list the names of the six students they spend the most time with. The data may therefore be right censored: if the true number of contacts exceed the limit of six named students, the network will not represent all potential interactions. Moreover, single links may represent peers an individual would like to be friends with, rather than actual contacts. However, similar contact groupings can be observed in the networks with single and mutual links (Figure 3), suggesting that consistent contact patterns were being captured. In self-reported questionnaires there is also the issue of recall bias [26], but it has been shown that children are capable of achieving similar levels of accuracy to adults for simple objective questions [18], and that links between school classes in networks inferred from surveys are consistent with networks measured using wearable sensors [36]. Although we worked closely with the schools to ensure questionnaires were conducted as consistently as possible, the times between rounds of data collection also fluctuated based on logistical constraints, such as term dates. Given that the ‘true’ contact network was never observed, only measured contacts in each round, our analysis of out-of-sample predictions reflects consistency between rounds, rather than comparison to a known baseline network. There was also a low response rate from some classes, mostly as a result of parents either not receiving, completing or returning a consent form. A one off survey may have been more effective in terms of response rate [23], but it would not have been possible to analyse the longitudinal patterns we were interested in. One option for future work would be to extend the data collection period over a year or two, rather than over five months we considered, to see whether the consistency in contact patterns held over multiple school years. Although this would be provide longer term insights, it would also potentially introduce further practical challenges both for researchers and school staff.

We focused on close social contacts in this study, as it has been suggested that these play a key role in transmission of respiratory infections [2, 41]. However, there may other important routes of transmission that are not captured by self-reported social mixing patterns, such as fomites and aerosols [42, 43]. An outstanding challenge for such infections is how to measure social contacts at the time of infection, and hence link transmission events with the interactions that generated them. Bacterial infections, for which there can be high rates of nasopharyngeal carriage, may therefore be a good target for future research into this question [41].

## Acknowledgements

We wish to thank the year 10 and year 7 students, and staff, at the schools that made this research possible: St Bonaventure’s, London; Highgate School, London; St Pauls, Burgess Hill; and one other school. We would also like to thank the Millenium Mathematics Project for helping with recruitment of schools and Julia Carney for assistance with data entry. This project was supported by a Wellcome Trust People Award (105107/Z/14/Z). AJK was supported by a Sir Henry Dale Fellowship jointly funded by the Wellcome Trust and the Royal Society (grant Number 206250/Z/17/Z) and the Medical Research Council (MR/K021524/1).

---

**Table S1:**
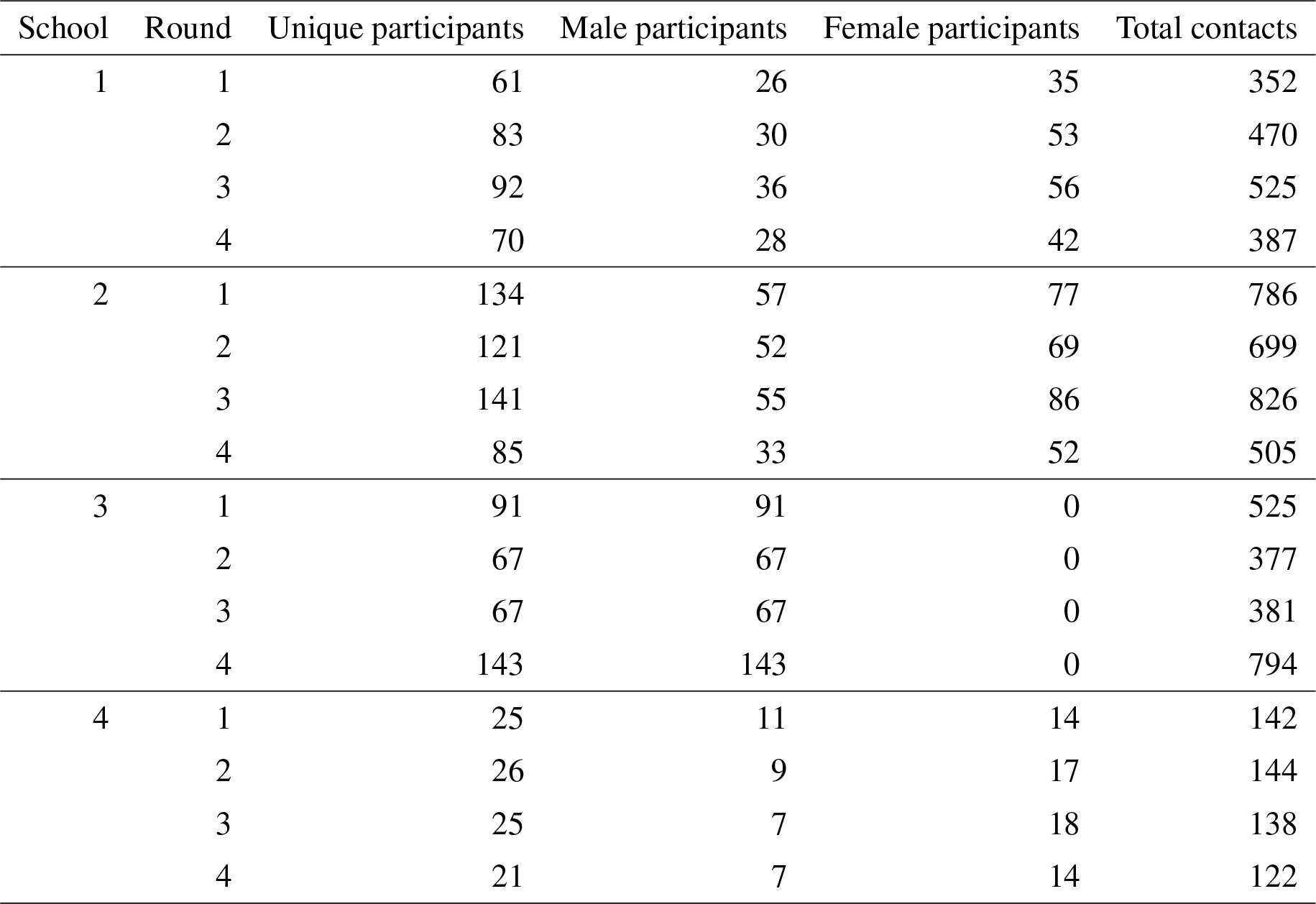
Summary of survey data from the four schools in each round of data collection (Table 1).

